# Perilipin 5 Phosphorylation is Dispensable for Upregulation of Hepatic Lipid Metabolism Genes upon Fasting but Required for Insulin Receptor Substrate 2 Expression in Male Mice

**DOI:** 10.1101/2024.11.09.622792

**Authors:** Corinne E Bovee, Ryan P Grandgenett, Michelle Trevino, Sucharita Dutta, Spencer J Peachee, Shayla Kopriva, Farakh Haider, Siming Liu, Gourav Bhardwaj, Christie Penniman, Brian T O’Neill, Yumi Imai

## Abstract

**Objective:** Perilipin 5 (PLIN5) is a lipid droplet protein highly expressed in cells that actively oxidize fatty acids. Previous in vitro studies have revealed that PLIN5 phosphorylation (p-PLIN5) at serine 155 by PKA is critical for transcriptional regulation of PPARa target genes by which PLIN5 adapt cells for fatty acid oxidation. We aim to determine the extent of p-PLIN5 in vivo and the consequence of impaired PLIN5 phosphorylation in the liver by using a whole-body knock-in of phosphorylation resistant PLIN5 (SA/SA) in mice.

**Methods:** We measured PLIN5 and p-PLIN5 with mass spectrometry and Phos-tag gels. We assessed serum chemistry in WT and SA/SA mice upon fasting. RNA sequencing and qPCR compared the gene expression in the liver of SA/SA and WT mice after overnight fast.

**Results:** Plin5 phosphorylation at S155 was increased in the liver LD fraction of fasted mice compared with that of fed mice by mass spectrometry (p<0.05). qPCR of key lipid metabolism genes did not differ between WT and SA/SA liver upon fasting. Male SA/SA mice had a higher fasting blood glucose (p<0.05) without a difference in body weight, serum insulin, or serum lipids. IRS2 was reduced in the liver of fasted male SA/SA mice (p<0.05).

**Conclusion:** PLIN5 S155 phosphorylation is dispensable for the upregulation of lipid metabolism genes important for fasting response in vivo. Impaired phosphorylation also had little effect on serum lipids or liver TG. However, SA/SA mice showed decreased IRS2 expression in the liver, which may contribute to glucose intolerance in SA/SA male mice.

## 1. Introduction

The lipid droplet (LD) is an organelle that serves as a hub for intracellular lipid metabolism with its ability to store neutral lipids and release lipids in a temporally and spatially regulated manner [1]. Five members of the perilipin family of protein (PLIN1-5) reside on the surface of LDs and confer the regulation on lipid metabolism, each with a unique expression pattern and molecular characteristics [2; 3]. PLIN1 is the best studied PLIN whose expression is restricted to adipocytes and adrenocortical cells. PLIN1 allows massive accumulation of triglycerides (TG) in adipocytes at the basal status. Upon activation of protein kinase A (PKA), such as during fasting, adipocyte PLIN1 facilitates rapid degradation of TG by lipolysis to release fatty acids (FA) into circulation for the use in extra adipose tissues [3]. Perilipin 5 (PLIN5) is highly expressed in cells and conditions that heavily depend on fatty acid oxidation (FAO) such as brown adipocytes, cardiomyocytes, oxidative skeletal muscle cells, hepatocytes upon fasting, and pancreatic islets upon fasting [4; 5]. Studies primarily performed in cell culture models have revealed that PLIN5 has molecular characteristics beneficial for adaptation to high levels of FAO. At the basal status, PLIN5 prevents lipolysis by sequestrating adipose triglyceride lipase (PNPLA2) from its co-activator ABHD5, thus expanding the TG pool [6–8]. Upon PKA activation, PLIN5 promotes lipolysis by PNPLA2 in part by increasing the interaction between PLIN5 and PNPLA2 [9–11]. Mouse PLIN5 contains one PKA consensus sequence (RRXS) at amino acid S155 that is RRWS [12], which is considered to be necessary to upregulate lipolysis under PKA activation [10]. In addition, PLIN5 phosphorylation at S155 by PKA triggers nuclear translocation of PLIN5 that is proposed to transport monounsaturated FA to nuclei and to activate peroxisome proliferator-activated receptor-gamma coactivator (PGC1a) through deacetylation catalyzed by Sirtuin 1 (SIRT1), collectively upregulating the expression of PPARa target genes important for FAO, mitochondrial mass/function, autophagy, and inflammation [7; 8; 11; 13; 14]. Importance of lipolysis in the regulation of PPARa target genes is also demonstrated in mouse models in which *Pnpla2* downregulation in cardiomyocytes and hepatocytes reduced the expression of PPARa target genes [15; 16]. The transcriptional change mediated by lipolysis and PLIN5 is considered to adapt cells to the increased demand of FAO and stress associated with FAO. Thus, PKA responsiveness of PLIN5 appears to be critical for upregulation of lipolysis and the transcriptional regulation in cells under high demand of FAO. However, the impact of PLIN5 phosphorylation in vivo remains less defined since most studies of PLIN5 phosphorylation utilized in vitro models with pharmacological stimulation of PKA [9; 13; 17]. Two studies reported phenotypes of mice expressing phosphorylation-resistant PLIN5 in cardiomyocytes and hepatocytes, but both compared WT and phosphorylation-resistant PLIN5 overexpressed under exogenous promoters and the impact of PLIN5 phosphorylation on hepatic PPARa gene expression was not directly compared between WT and phosphorylation resistant PLIN5 expressed at physiological levels [8; 11]. Here, we address the extent of PLIN5 phosphorylation in vivo and the consequence of impaired phosphorylation using a model in which phosphorylation-resistant PLIN5 was expressed under the endogenous promoter (SA/SA mice). The liver upon fasting was tested as a physiological condition when PPARa activation plays a critical role in switching lipid metabolism to increase FAO [18]. Also, PKA-dependent PLIN5 phosphorylation in the liver is expected to increase during fasting due to the rise of cAMP [19; 20]. We demonstrated that the total PLIN5 protein level is increased during fasting in the liver with the proportional increase of phosphorylated PLIN5. SA/SA knock-in mice that replaced S155 PLIN5 with alanine showed significant reduction in PLIN5 phosphorylation in the liver upon fasting supporting that S155 is the major phosphorylation site of PLIN5. However, liver TG, serum lipids, serum ketone, and the expression of hepatic lipid metabolism genes that are known to be regulated by PPARa during fasting were not altered in SA/SA mice upon fasting. However, old male SA/SA mice showed mild glucose intolerance that was associated with reduced expression of IRS2 in the liver.

## 2. Methods

### 2.1. Cell culture

Adenovirus expressing wild type (WT) mouse PLIN5 (Ad-PLIN5) and PLIN5 in which serine 155 was mutated to alanine (SA) were created by Vector Biolabs (Philadelphia, PA) with confirmation of cDNA sequence. Hepatocytes from C57BL/6J mice were provided by Triangle Research Labs (Charlottesville, VA). Hepatocytes were transduced in suspension with 100 plaque forming units (pfu) per cell of Ad-PLIN5 for 1 h in serum-free DMEM containing 22.2 mM glucose, plated at 2.5×10^5^ cells/well in a 6-well plate, and cultured overnight in the presence of 10% FBS DMEM at 37 °C in 5% CO_2_. Then, Ad-PLIN5 transduced hepatocytes were incubated for 24 h in DMEM containing 9 mM glucose plus 0.5 mM oleic acid (OA) conjugated to 1% FA free albumin to promote LD formation. On the next day, cells were incubated in serum-free 9 mM glucose DMEM for 5 h, treated with and without 20 μM H89 dihydrochloride (Cell Signaling, Beverly, MA) for 30 min followed by incubation with 1 mM 8-bromoadenosine 3’,5’-cyclic monophosphate (8-Br-cAMP, Sigma, St Louis, MO) for 15 min, washed with cold PBS, snap frozen by submerging wells in liquid nitrogen and scraping into cold IP buffer (249 μM mannitol, 0.5% triton-X, 50 mM Tris HCl pH 7.5 supplemented with protease and phosphatase inhibitor cocktails (Sigma)) for further analysis. AML12 cells (CRL-2254, American Tissue Culture Collection, Manassas, VA) were maintained in DMEM:F12 medium supplemented with 10% FBS, 10 μg/ml insulin, 5.5 μg/ml transferrin, 5 ng/ml selenium, and 40 ng/ml dexamethasone. AML12 cells were transduced with Ad-PLIN5 or Ad-SA PLIN5 as in hepatocytes, loaded with 0.2 mM OA starting from the next day, and harvested on the third day as for hepatocytes after 1 h incubation with or without 8-Br-cAMP.

### 2.2. Animal studies

Experiments were performed in accordance with the Institutional Animal Care and Use Committee guidelines of University of Iowa. Mice were housed 5/cage in 12 h light-dark cycle at 22 °C, allowed free access to water, and fed regular rodent chow (7319 Teklad global diet) except during fasting experiments. Female leptin mutant *ob/ob* mice and WT mice (BL6J) in C57BL/6J background were obtained from Jackson Laboratory (Bar Harbor, ME). For some experiments, male WT C57BL/6NJ (BL6NJ) mice from Jackson Laboratory were used. Treadmill exercise test was performed as previously described [21]. In brief, mice after acclimation were placed on the treadmills at a 10° angle and ran at the starting speed of 4 m/min with 2 m/min increase in the speed every 10 min for 70 min. After 70 min, speed was increased by 4 m/min every 10 min until exhaustion, determined by the mice being unable to leave the treadmill shock grids for 10 consecutive seconds. Shock grids were set to irritate but not harm the mice. Mice in which serine at 155 (AGT) of PLIN5 was replaced to alanine (GCC) by CRISPR-Cas9 gene editing were created at Genome Editing Core Facility at University of Iowa (SA/SA mice). The correct replacement of targeted sequence was confirmed by DNA sequencing in a founder mouse. Genotype of subsequent offsprings was performed as in supplementary methods. At the time of harvest, the liver and the heart were snap frozen in liquid nitrogen and stored at −80 °C until analyses.

### 2.3. Lipid droplet semi-purification

LD were semi-purified by a single centrifugation protocol published by Harris *et al* [22] with the following modifications (Fig. 1A). 400-600 mg of the left lobe of liver snap frozen upon harvest was transferred to ice-cold 60% sucrose lysis buffer [22], crushed with pestle, and incubated on ice for 10 min with vortex every 5 min. After adding lysis buffer without sucrose for a final volume of 1 ml, liver samples were incubated on ice for additional 10 min with vortex every 5 min. Liver samples were then homogenized with a hand held homogenizer and incubated on ice for 15 min with vortex every 5 min. 300 μl of lysis buffer with blue food color dye was layered on top of homogenate and centrifuged at 20,000 x *g* for 2.5 h at 4 °C, which enriches LD in a floating layer on top of the blue dye layer (Fig. 1A2). LD layer was transferred to a new tube for further analysis.

**Figure 1.**
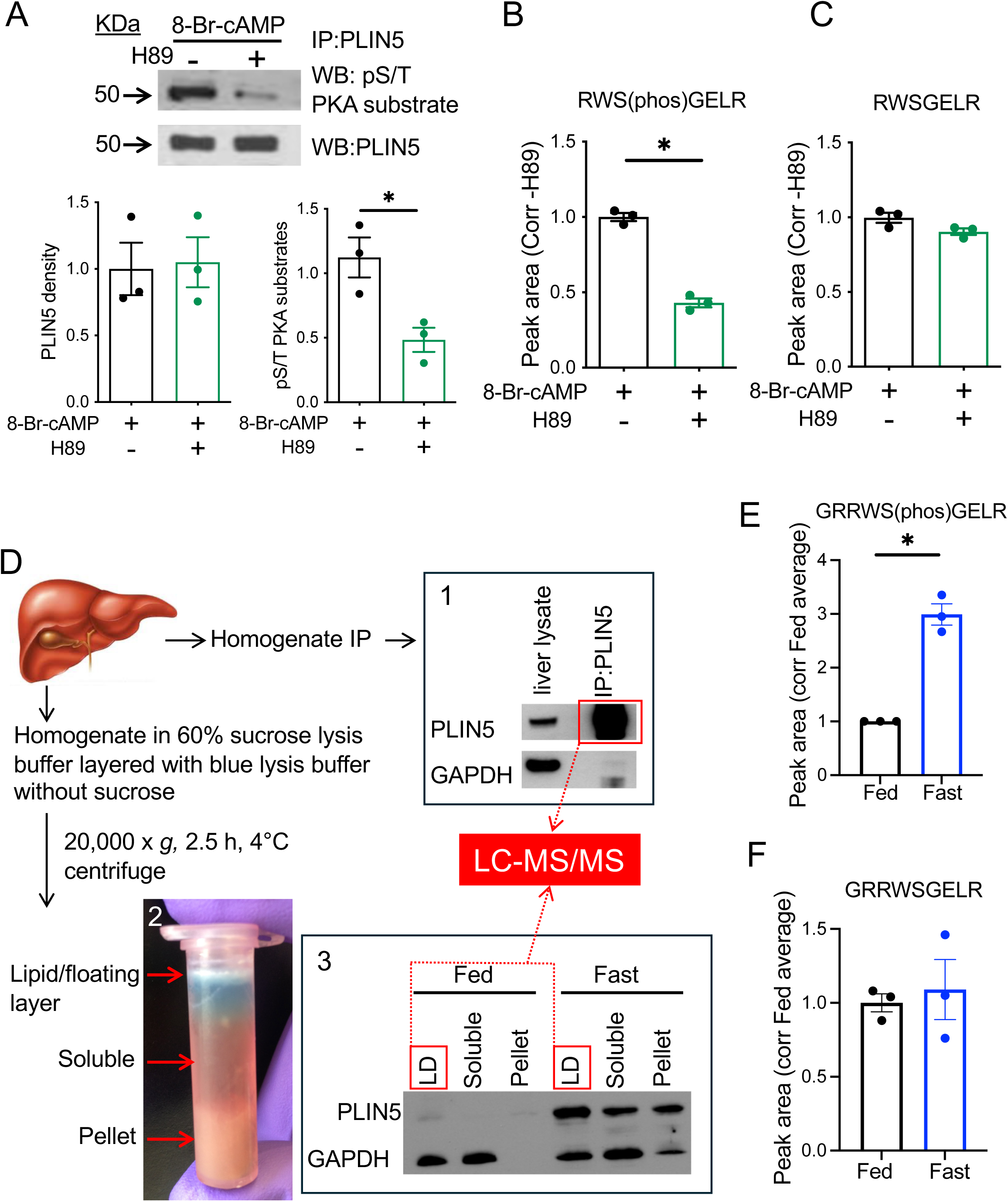
S155 PLIN5 phosphorylation detected by mass spectrometry is increased in the LD fraction of liver from fasted mice. (A) Western blot (WB) of PLIN5 immunoprecipitated (IP) from hepatocyte lysate overexpressing wild type (WT) PLIN5 and treated with 8-Br-cAMP ± H89 was performed for indicated antibodies. Representative images and densitometry data are shown. (B, C) Spectral peak areas of PLIN5 peptides RWS(phos)GELR (B) and RWSGELR (C) were obtained by liquid chromatography tandem mass spectrometry (LC-MS/MS) for tryptic peptides of (A). (D) Schematic diagram of PLIN5 enrichment from WT mouse liver homogenate and LC-MS/MS analysis. 1. WB of PLIN5 and GAPDH comparing whole liver lysate and PLIN5 IP from the fast liver. 2. Photo depicts the separation of lipid/floating layer, soluble layer, and pellet after the single centrifugation step as in Methods. 3. WB compared PLIN5 and GAPDH levels in the three fractioned layers from the livers of fed and fasted mice. Fractions applied to LC-MS/MS are indicated by red. (E, F) Spectral peak areas for PLIN5 peptides GRRW(phos)SGELR (E) and GRRWSGELR (F) obtained from 67 mcg protein of fed and fast liver LD fractions in 3. Data are means ± SEM; n = 3. \**p* < 0.05 by Student’s t test.

### 2.4. Immunoprecipitation and Western blotting

For immunoprecipitation (IP), tissues and cells were lysed in IP buffer. Protein concentrations were measured with Bio-Rad DC Protein Assay (Bio-Rad, Hercules, CA). PLIN5 IP used guinea pig anti-PLIN5 antibody (Progen Biotechnik, Heidelberg, Germany) and Pierce’s Classic Immunoprecipitation Kit (Thermo Scientific, Waltham, MA) according to the manufacture’s instruction. Western blots were performed using 20 to 40 μg/lane of protein and detected by chemiluminescence or Odyssey CLx imaging system (Licor, Lincoln, NE). Densitometric analyses were performed with Image J. Antibodies and dilution used are in supplementary table 1. For PLIN5 in-gel digestion, 67 mcg of protein from LD fraction was run on a Bis-Tris gel. For separation of p-PLIN5, SuperSep Phos-tag (50 µmol/L) 12.5% precast gel (Wako Chemical, Richmond, VA) was used. 12-15 μg of liver lysate per lane was run on the gel and the gel was washed prior to transfer following manufacturer’s instructions. Proteins were transferred onto a nitrocellulose membrane and blocked in PBS containing 0.1% tween 20 and 3% BSA prior to antibody treatment.

### 2.5. PLIN5 in-gel reduction, alkylation, denaturation and trypsin digestion

The gel stained with Page Blue was de-stained and sequentially washed with buffers containing 50 mM ammonium bicarbonate, 50% acetonitrile, and 80% acetonitrile. The bound proteins were reduced with 1 ml of 40 mM dithiothreitol for 25 min at 56 °C. The gels were rinsed with 1 ml of 50 mM ammonium bicarbonate buffer and the reduced proteins were alkylated with 1 ml of 50 mM iodoacetamide for 30 min at 25 °C in the dark with constant mixing. The iodoacetamide was discarded and the gel bound proteins were digested with 0.5 ml of 20 ng/µl trypsin (Promega, Madison, WI) in 50 mM ammonium bicarbonate buffer at 37 °C with constant mixing for 12 h. After digestion, the tryptic fraction released into 50 mM ammonium bicarbonate buffer was collected, and the gels were washed with 50 mM ammonium bicarbonate to collect any remaining tryptic peptides. The ammonium bicarbonate buffer containing the tryptic peptides was dried using a Speed-Vac apparatus and stored at 4 °C prior to mass spectrometric analysis. The dried samples were dissolved with 20 µl of 0.1% formic acid in water. 2 µl each of sample was analyzed by Liquid Chromatography Tandem Mass Spectrometry (LC-MS/MS) using a Q-Exactive mass spectrometer (Thermo Fisher) with an Easy NanoLC-1000 system as in supplementary methods,

### 2.6. Colorimetric assays

Saphenous blood or cardiac blood collected at harvest was centrifuged at 2,000 x g for 10 min at room temperature to obtain serum. TG (TR22421 from ThermoFisher), β-hydroxybutyrate (Stanbio, Boerne, TX), non-esterified FA (Wako Chemicals), and mouse insulin (Alpco, Salem, NH) were measured according to manufacturers’ protocols. Liver TG was measured using an aforementioned TG colorimetric assay after Folch extraction as we published and values were corrected for protein contents [23].

### 2.7. mRNA and quantitative PCR

Total RNA from liver tissues (50-100 mg) were prepared using Qiagen RNeasy kit (Qiagen, Valencia, CA) according to the manufacture’s protocol and cDNA was synthesized as published [24]. Gene expression was assessed as previously described [24] using ABI TaqMan commercial primers (Applied Biosystems). Results were expressed using mouse PPIB as a reference gene.

### 2.8. Bulk RNA-sequence of the liver

RNA was extracted from the liver using TRIzol Reagent (Invitrogen, Carlsbad, CA) followed by RNA Clean & Concentrator-5 (Zymo Research, Irvine, CA) kit with DNase I treatment according to manufacturer instructions. An RNA integrity number (RIN) was measured by RNA ScreenTape (Agilent Technologies, Santa Clara, CA) and RNA with RIN above 7.2 was used. Gene expression profiling by bulk RNA-Seq was performed by the University of Iowa Genomics Division as in supplementary methods. The raw FASTQ files and associated metadata have been made available for download at GEO accession GSE275608.

### 2.9. Glucose homeostasis

Glucose tolerance test (GTT) was done after overnight fasting by loading 1 mg/g BW glucose intraperitoneally (i.p.) and tail blood glucose was measured at indicated time using a handheld glucometer. Insulin tolerance test (ITT) was performed after 6 h fasting by 0.75 mU/g BW regular insulin i.p. and glucose was measured at in GTT.

### 2.10. Statistical analysis

Data are presented as mean ± standard error of mean (SEM) unless otherwise stated in the figure legends. Differences of numeric parameters between two groups were assessed with Student’s t-tests. Welch correction was applied when variances between two groups were significantly different by F test using Prism 10 (GraphPad, La Jolla, CA). Multiple group comparisons used one-way ANOVA with post hoc as indicated. When two categorical variants exist, two-way ANOVA test was performed. A p < 0.05 was considered significant.

## 3. Results

### 3.1. Fasting increases PLIN5 phosphorylated at S155 in the mouse liver

While S155 has been shown to be phosphorylated in a PKA dependent manner in vitro, phosphorylation of PLIN5 at S155 in response to physiological stimuli has not been shown in the liver [11; 13]. To establish a protocol to detect S155 PLIN5 by LC-MS/MS, primary hepatocytes overexpressing PLIN5 were treated acutely with cAMP analog, 8-Br-cAMP in the presence or absence of a PKA inhibitor H89. PLIN5 was enriched by IP and analyzed by Western blot (Fig. 1A) and LC-MS/MS (Fig. 1B and C) in parallel. In Western blot, H89 markedly reduced phospho-PKA substrate signal of PLIN5 IP in support of PKA dependency of PLIN5 phosphorylation (Fig. 1A). LC-MS/MS of these IP samples yielded RWS(phos)GELR as a dominant tryptic peptide containing S155 of PLIN5 in primary hepatocytes (Fig. 1B, Supplementary Fig. 1A). While abundance of RWSGELR peptide was similar in both groups, RWS(phos)GELR peptide was reduced in H89 treated hepatocytes showing good agreement with Western blot (Fig. 1B-C). We then analyzed PLIN5 enriched by IP of the liver lysate from male WT BL/6NJ mice fasted overnight by LC-MS/MS (Fig. 1D1). Fasting caused 11% weight loss, 51% reduction of blood glucose, 701% increase of serum beta-hydroxybutyrate, and 225% increase of PLIN5 protein by Western blot of liver lysate as expected (supplementary Fig. 1B-E) [4]. Tryptic peptides of PLIN5 IP samples from the fasted mouse liver analyzed by LC-MS/MS showed that GRRWS(phos)GELR is the major peptide that includes S155 in the liver. Then, we moved to the quantitative comparison of GRRWS(phos)GELR peptide in the fast and fed liver. We chose to utilize LD fraction semi-purified from liver homogenate due to significant difference in total PLIN5 protein levels that may not allow the proportional recovery of PLIN5 after IP (Fig. 1D2). PLIN5 was primarily detected in LD fraction in both fed and fast liver and was much more abundant in the fast liver (Fig. 1D3). When the equal amount of protein was loaded, the area of the spectral peak for GRRWS(phos)GELR peptide in LD fraction was significantly increased in the fast liver compared with fed liver (Fig. 1E), while that for GRRWSGELR did not differ (Fig. 1F). Thus, LC-MS/MS indicated that fasting increases phosphorylation of PLIN5 at S155 in the liver LD fraction of fasted mice.

### 3.2. Phos-tag gel allows quick separation of non-phosphorylated and phosphorylated PLIN5

While mass spectrometry provides measurement of p-PLIN5 specifically at S155, it is time and resource consuming. Phos-tag gel contains the Phos-tag molecule that specifically binds to phosphorylated proteins and slows their migration during SDS-page [25]. We tested whether Phos-tag gel can be utilized to differentiate non-phosphorylated vs phosphorylated forms of S155 PLIN5. First, AML12 cells expressing WT and SA PLIN5 were stimulated by 8-Br-cAMP and cell lysates were run on Phos-tag gel. As shown in Fig. 2A, WT PLIN5 showed predominance in the top band at baseline with disappearance of bottom band in the presence of 8-Br-cAMP. In contrast, SA PLIN5 showed predominance of the bottom band at the baseline that did not change in the presence of 8-Br-cAMP. Moreover, Phos-tag gel showed a stepwise reduction of p-PLIN5 in the liver lysate from PLIN5^SA/+^ (SA het) and PLIN5^SA/SA^ knock-in (SA/SA) male mice fasted overnight compared with PLIN5^+/+^ (WT) mice confirming that S155 is the major phosphorylation site of PLIN5 in the liver PLIN5 in vivo (Fig. 2B). Thus, Phos-tag gel can detect changes in PLIN5 phosphorylation at S155 both in cell culture and mouse models (Fig. 2A,B).

**Figure 2.**
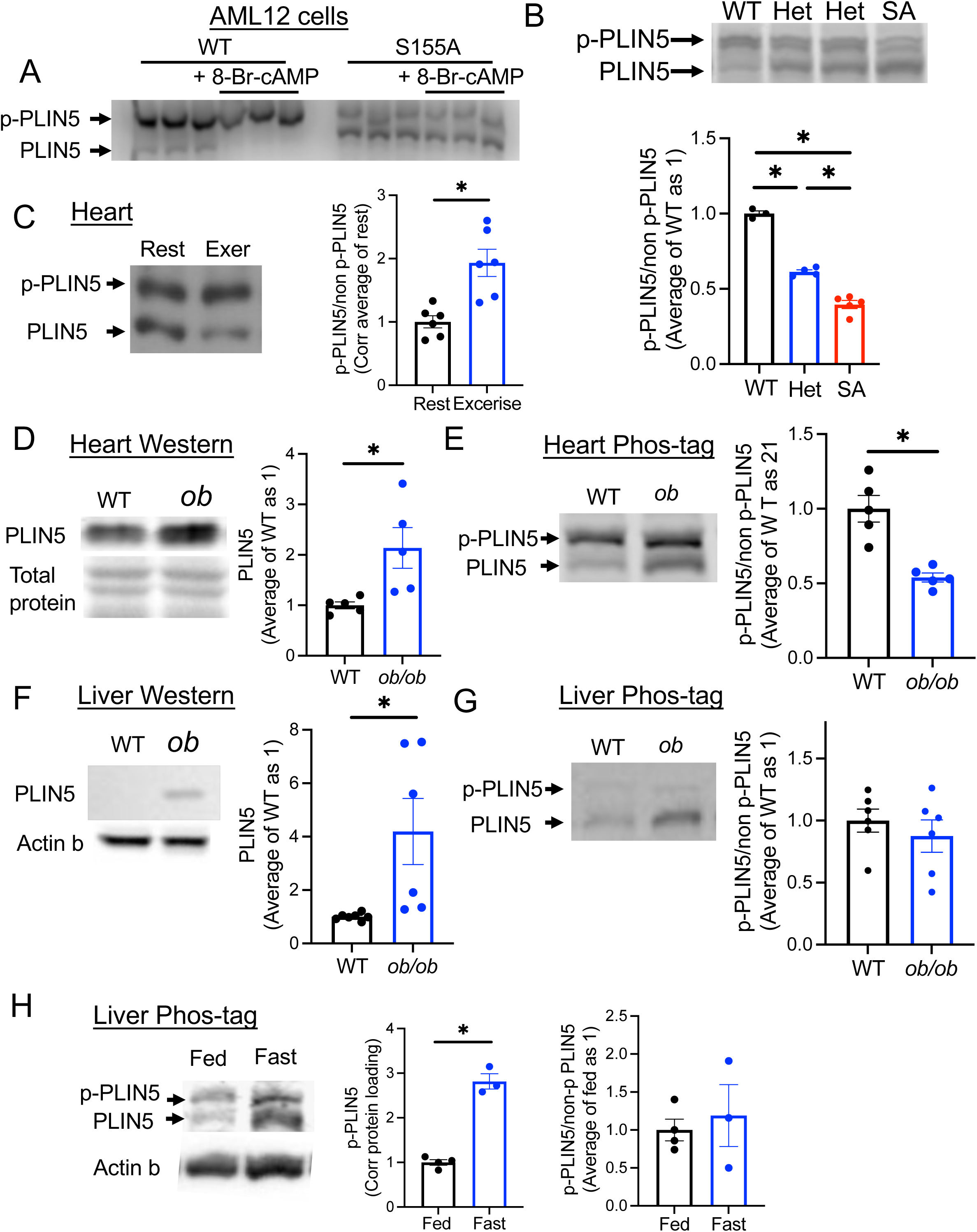
Phos-tag gel allows the detection of p-PLIN5. (A) Cell lysate from AML12 cell overexpressing wild type (WT) or S155A PLIN5 and treated with or without 8-Br-cAMP was run on Phos-tag gel and immunoblotted by anti-PLIN5 antibody. Representative image of 3 experiments. (B) Phos-tag gel of the liver from fasted 1-year-old male WT, SA het (het) and SA/SA (SA) mice assessed the ratio of phosphorylated (p-PLIN5) and non-phosphorylated (non p-PLIN5). n=3-5. (C) Phos-tag gel assessed p-PLIN5 and non p-PLIN5 in the heart of WT mice with (Exer) and without (Rest) treadmill exercise. n=6. (D-G) The heart (D, E) and the liver (F, G) lysates from female WT and *ob/ob* mice were analyzed for PLIN5 by regular Western blot (D, F) and Phos-tag gel (E, G), n=5-7. Western blot of PLIN5 was normalized by total protein staining for the heart (D) and actin b for the liver (F). (H) Phos-tag gel assessed the abundance of p-PLIN5 and non p-PLIN5 in the total liver lysate of fed or overnight fasted male mice. n=3-4. (B-H) Representative gels and densitometry data are shown. Data are mean ± SEM. (B) *p<0.05 by One-way ANOVA with Sidak’s multiple comparison test. (C-H) *p<0.05 by Student’s t test.

### 3.3. Phos-tag gel detects the change in PLIN5 phosphorylation in vivo in the liver and the heart

We then asked whether the proportion of p-PLIN5 can be altered in conditions that increase FAO or PLIN5 expression in vivo. During exercise, the heart activates sympathetic tone that increases intracellular cAMP and FAO [26]. The heart harvested from WT mice run on treadmill (run distance 1149 ± 100 m, mean ± SEM, n=6) showed increased proportion of p-PLIN5 (Fig. 2C). The validity of exercise was supported by the increase in blood lactate levels (supplementary Fig. 2A). Next, we tested whether the proportion of p-PLIN5 is altered when obesity increases PLIN5 protein in the liver and heart (Fig. 2D-G) using *ob/ob* mice (body weight and blood glucose in supplementary Fig. 2B, C). Western blot showed the increase of PLIN5 in both the liver and the heart of *ob/ob* mice compared with WT mice (Fig. 2D, F). The proportion of p-PLIN5 was reduced in the heart but not altered in the liver of *ob/ob* mice compared with WT mice (Fig. 2E, G). For the liver from fasted WT mice, p-PLIN5 corrected for protein was increased in Phos-tag gel (Fig. 2H). However, there was not statistically significant change in the proportion of p-PLIN5 as non-phosphorylated PLIN5 in total liver lysate was also increased upon fasting (Fig. 2H). Combined with LC-MS/MS data (Fig. 1E, F), p-PLIN5 is increased during fasting especially in LD in the liver, but this may be proportional to the increase in total PLIN5 contents in the liver based on Fig. 2H. Also, Phos-tag gel revealed that the extent of p-PLIN5 varies depending on tissues and context with more dynamic changes in the heart than in the liver.

### 3.4. Parameters of lipid metabolism and the expression of hepatic PPARa target genes important for fasting response are not different between WT and SA/SA mice

One proposed consequence of p-PLIN5 is upregulation of PPARa target genes by increasing lipolysis and activating Sirt1/PGC1a pathway [13; 14]. PPARa is the master transcriptional regulator mediating dramatic changes in the expression of lipid metabolism genes during fasting in the liver [18]. Previously, a subset of PPARa target lipid metabolism gene was shown to be transcriptionally regulated by lipolysis and/or PLIN5 phosphorylation in cultured cells [13; 14]. Thus, we used SA/SA mice to assess whether S155 PLIN5 phosphorylation is necessary for expression of PPARa target genes during fasting in the liver. Body weight, serum TG, serum non-esterified FA (NEFA), and serum beta-hydroxybutyrate did not differ between WT, het, and SA/SA knock in mice after fasting in 1-year-old male mice (Fig. 3A-D) or 4-month-old female mice (supplementary Fig. 3A-D). Then, we chose *Pgc1a, Ppara, Cpt1a, Acot1,* and *Plin5* as lipid metabolism genes of interest as their expression was previously shown to be regulated by p-PLIN5 [13; 14]. While the expression of *Ppara* can be regulated by glucocorticoid and *Cpt1a* by CREB3L3 during fasting in the liver [18], *Acot1,* and *Plin5* are regulated by PPARa [27; 28]. The expression of *Ppara, Cpt1a, Acot1, and Plin5* was markedly upregulated after overnight fasting in the liver of male WT mice as expected (Fig. 3E). Surprisingly, there was little difference in the expression of these genes between WT and SA mouse liver after fasting in either males or females (Fig. 3F, G). Liver TG contents were not altered in fasted male and female SA/SA mice either (Fig. 3H-I).

**Figure. 3.**
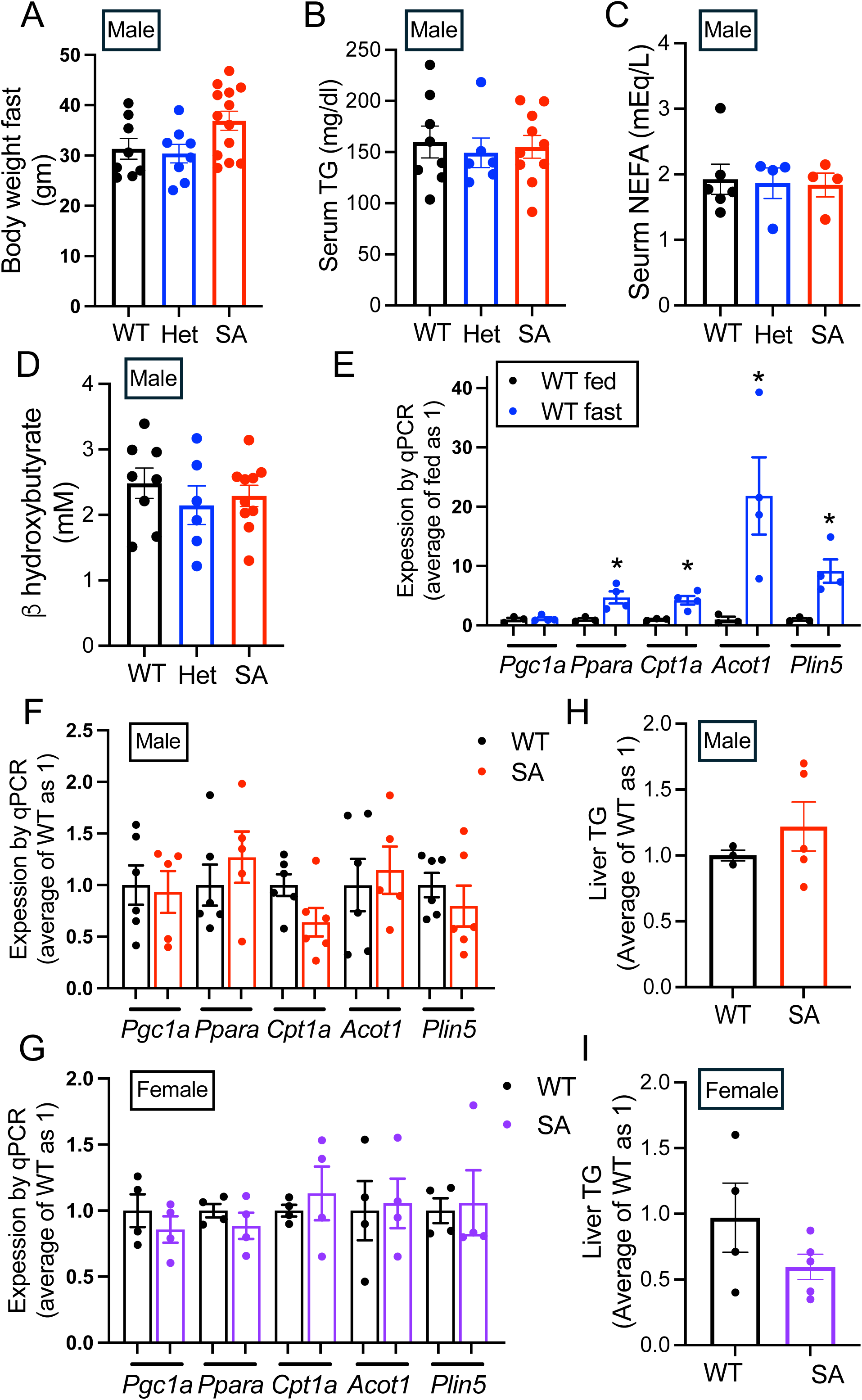
Serum chemistry and hepatic lipid metabolism gene expression in fasted PLIN5 SA/SA mice. (A) Body weight, (B) serum TG, (C) serum non-esterified fatty acids (NEFA), and (D) serum β-hydroxybutyrate of 1-year-old male mice after overnight fast. Each dot represents one mouse. n= 6-8 (WT), 4-8 (het), and 4-13 (SA). (E-G) qPCR compared the expression of genes in lipid metabolism in the liver of (E) WT mice fasted overnight (n=3) vs fed *ad libitum* (n=4), (F) 1-year-old WT (n=6) vs SA mice fasted overnight (n=5-6), and (G) 6-month-old WT (n=4) vs SA female mice fasted overnight (n=4). (H, I) TG contents in the liver corrected for protein contents were obtained in (H) male and (I) female mice after overnight fast and expressed taking average of WT liver as 1. n= 3 (WT) and 5 (SA) for males, and 4 (WT) and 5 (SA) for females. Data are mean ± SEM. *p<0.05 by Student’s t test.

### 3.5. RNA sequencing of female SA/SA liver showed limited number of differentially expressed genes

With little change in parameters of lipid metabolism including blood chemistry, liver TG levels, and lipid metabolism gene expression, we performed unbiased analysis of liver gene expression by RNA sequencing to identify genes differentially regulated by the loss of S155 phosphorylation. Female mouse liver was used considering that fasting response is reported to be more pronounced in females [29]. However, RNA sequencing revealed very small number of genes that are differentially expressed (Fig. 4A, supplementary table 2) with the notable reduction of *Irs1.* The reduction of *Irs1* was confirmed by qPCR in female SA/SA liver, but not seen in male SA/SA liver (Fig. 4B, C). Instead, *Irs2* was reduced in SA/SA male liver but not in female SA/SA liver by qPCR (Fig. 4B, C). In the liver of WT male mice, both *Irs1* and *Irs2* were highly upregulated by fasting (Fig. 4D). Western blot confirmed IRS2 is reduced in SA/SA male liver (Fig. 4E). However, the reduction of IRS1 was not demonstrated in SA female liver (Supplementary Fig. 3E). With changes in the expression of *Irs,* we compared glucose tolerance between WT and SA/SA mice. There was not significant difference in glucose tolerance between 4-month-old WT, SA het, and SA/SA male or female mice (supplementary Fig. 4A, B). However, 1-year-old SA/SA male mice were mildly glucose intolerant and hyperglycemic after overnight fasting (Fig. 5A-C). Insulin tolerance did not detect clear difference between WT and SA male mice either at young or old age (Fig. 5D, supplementary Fig. 4C). While some male mice in SA group showed elevated fasting serum insulin, the values were not statistical significance between three groups of mice (Fig. 5E). There was not significant difference in fasting glucose or serum insulin levels in female mice (supplementary Fig. 4D, E). With the increase in fasting blood glucose in 1-year-old male SA/SA mice, we tested the expression of *Pck1* and *G6PC*, genes that regulates gluconeogenesis and upregulated in WT liver after fasting (Fig. 5F). *G6PC* showed trend of reduction in male SA/SA liver (Fig. 5G), but no changes were seen in female SA/SA liver (supplementary Fig. 4F).

**Fig. 4.**
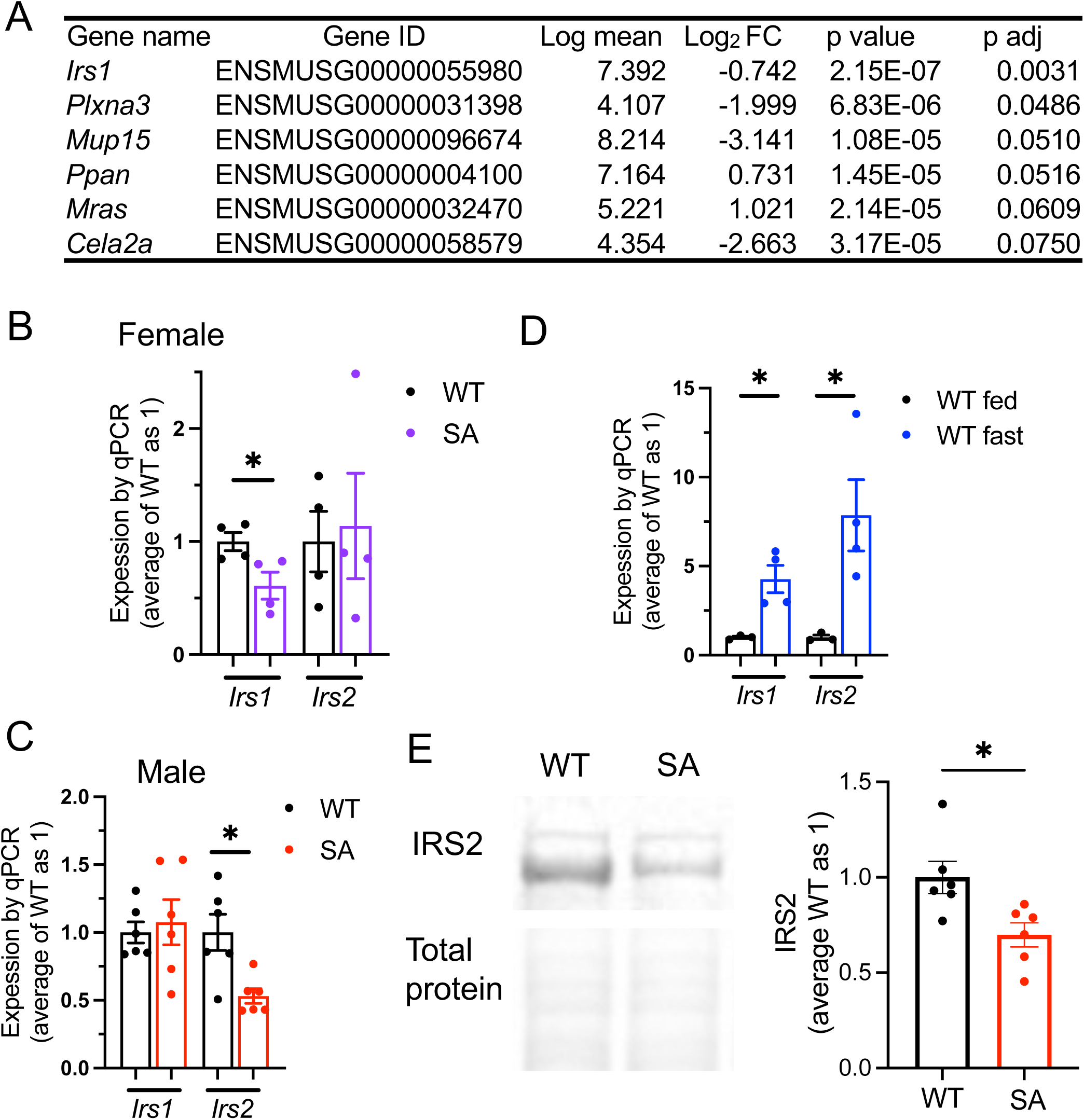
RNA sequencing of SA/SA liver identified differential expression of *Irs1* and *Irs2* in the liver. (A) The list of genes differentially expressed in the liver of female SA/SA (SA) mice compared with WT mice in RNA sequencing. FC; fold change. (B-D) qPCR compared the expression of *Irs1* and *Irs2* in the liver of (B) overnight fasted WT and SA female mice, n= 4 (C) overnight fasted WT and SA male mice, n= 6, and (D) overnight fasted and *ad libitum* fed WT male mice, n=3-4. (E) Western blot compared IRS2 and total protein levels in the livers of fasted WT and SA male mice. n=6. Data are mean ± SEM. *p<0.05 by Student’s t test.

**Figure. 5.**
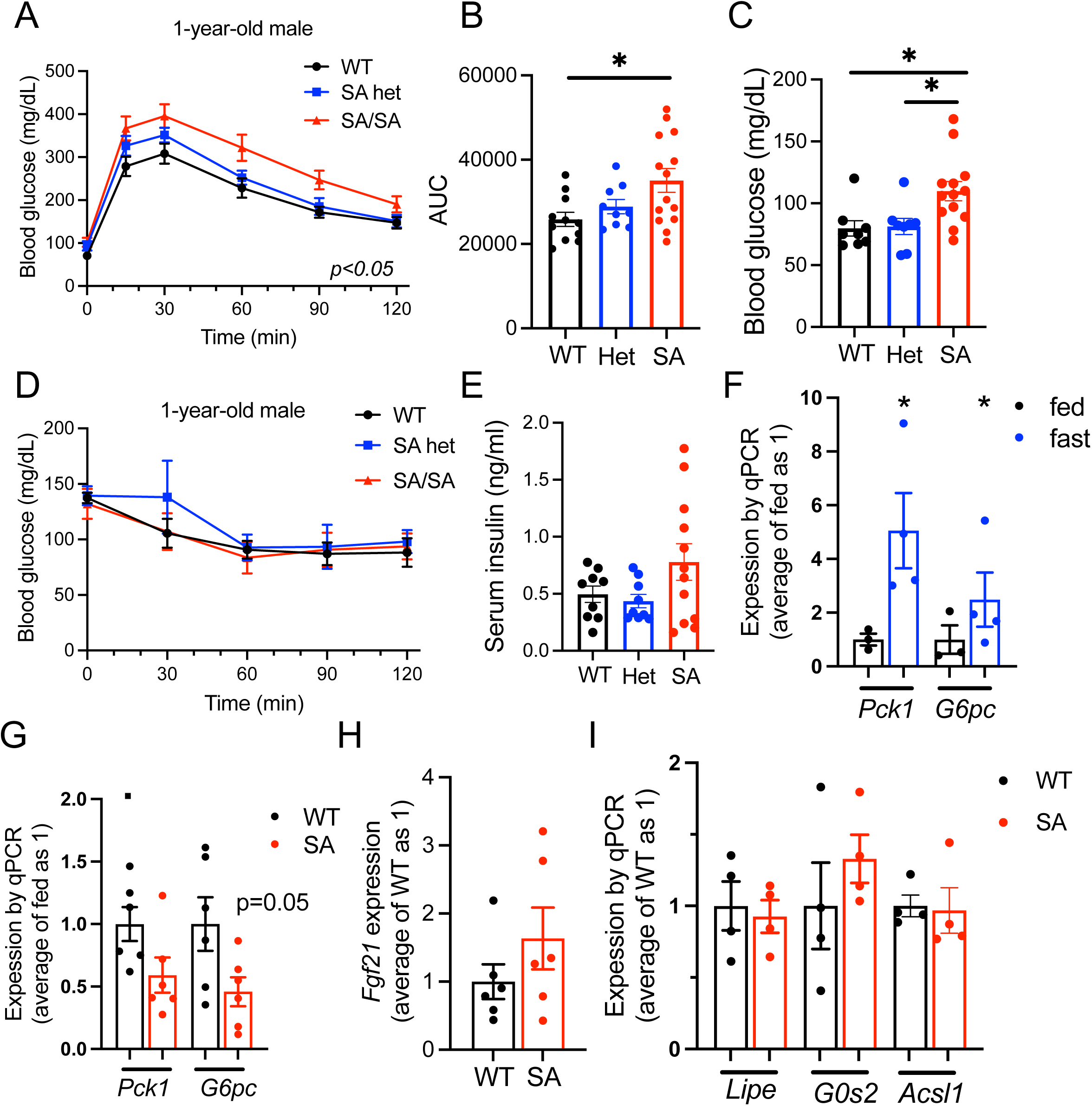
Glucose homeostasis and the expression of gluconeogenesis genes in the liver of PLIN5 SA/SA mice. (A) i.p. glucose tolerance test compared male WT, SA het (het), and SA/SA (SA) mice at 1-year-old. (B) Area under the curb (AUC) of (A). (C) Fasting glucose of (A). (D) i.p insulin tolerance test compared male WT, het, and SA mice at 1-year-old. (E) Serum insulin of 1-year-old male mice after overnight fasting. (A-E) n= 7-11 (WT), 3-9 (het), and 5-14 (SA). (F, G) qPCR compared the expression of *Pck1* and *G6pc* (F) in the liver of WT mice fasted overnight vs fed *ad libitum,* n=3-4 and (G) 1-year-old WT vs SA male mice fasted overnight, n=5-6. (H, I) qPCR compared the expression of (H) *Fgf21* and (I) lipid metabolism genes in the liver of 1-year-old WT vs SA mice fasted overnight. n=4-6. Data are mean ± SEM. (A) Two-way ANOVA test. (B, C) *p<0.05 by One-way ANOVA with Sidak’s multiple comparison test. (F, G) *p<0.05 by Student’s t test.

### 3.6. Male SA/SA liver did not show difference in Fgf21 expression, OXPHOS proteins, or inflammatory markers

As RNA sequencing was performed in females, we probed additional genes reported to be regulated by PLIN5 in other tissues and cultured cells. *Fgf21*is induced in the liver by fasting in PPARa dependent manner [30] and its increase is reported when PLIN5 is overexpressed in the heart and skeletal muscle [31; 32]. However, *Fgf21* expression was not altered in the liver of SA/SA male mice compared with WT mice upon fasting (Fig. 5H). The expression of *Lipe* and *G0s2* was reduced in the heart of mice overexpressing SA PLIN5 compared with mice without overexpression [8]. *Acsl1* is a PPARa target gene whose expression is reduced in the liver treated with PLIN5 antisense oligonucleotide (ASO) [14]. None of these genes were altered in the liver of fasted SA/SA male mice (Fig. 5I). Mitochondrial mass/function and inflammation are other pathways reported to be regulated by PLIN5 and its phosphorylation [13; 14]. There was no difference in OXPHOS proteins (supplementary Fig. 5A, B) or expression of inflammatory genes *Il-1b* and *Ccl2* in the liver of fasted SA/SA male mice (supplementary Fig. 5C). Collectively, the loss of S155 PLIN5 phosphorylation had limited impact on lipid metabolism, mitochondrial mass, or inflammation associated with fasting in the liver except for the reduced expression of IRS2 in male mice.

## 4. Discussion

The major finding of the current study is that S155 PLIN5 phosphorylation is largely dispensable for the upregulation of lipid metabolism genes that are considered to play an important role for adapting lipid metabolism upon fasting in the liver. During fasting, the liver receives NEFA released by adipocyte lipolysis and changes lipid metabolism drastically; the liver increases LD formation, upregulates FAO, and produces ketones. These changes heavily depend on a transcription factor PPARa [18]. Upon fasting, PPARa in the liver is activated by several mechanisms including the increased expression of PPARa, the activation of co-activators such as PGC1a, and better availability of NEFA [18]. PGC1a is an especially important co-activator for upregulation of FAO genes [18]. Interestingly, PLIN5 is reported to be transported to nuclei upon phosphorylation at S155 and increases transcription of PGC1a/PPARa genes by transporting FA to nuclei and/or increasing PGC1a activity through SIRT1 [13; 14]. In addition, PLIN5 phosphorylation has been shown to increase lipolysis, a process that supports PPARa activity providing an additional mechanism by which PLIN5 phosphorylation may support transcription of PPARa target genes [33]. Indeed, the expression of a number of PGC1a/PPARa target genes are shown to be regulated by PLIN5 phosphorylation in cultured cells; *CPT1a* (hepatocytes), *PGC1a* (hepatocytes, C2C12 myoblast), *PPARa* (hepatocytes) and *PLIN5* (C2C12 myoblast) [13; 14]. Thus, we assessed the contribution of PLIN5 phosphorylation in the liver upon fasting as a physiological condition in which PPARa target genes are highly upregulated and cAMP is increased [18; 19]. We confirmed that phosphorylated PLIN5 is increased in the liver upon fasting. However, genes proposed to be regulated through PGC1a/PPARa during fasting in the liver showed little changes in SA/SA mice. Targeted qPCR and unbiased RNA sequencing revealed little impact from the loss of S155 phosphorylation in the liver upon fasting except for differential gene expression of *Irs1* in females and *Irs2* in males.

Several plausible explanations exist for the limited impact of S155 PLIN5 mutation on gene expression in the fasted liver. First, the availability of FA as PPARa ligands might be maintained in SA/SA PLIN5 liver by mechanisms other than the transport of FA by p-PLIN5. While a detailed kinetic study is beyond the scope of the current study, we did not observe significant changes in liver TG indicating that there may not be a significant defect in lipolysis even when PLIN5 phosphorylation is prevented in vivo. Besides PLIN5, chaperon-mediated autophagy of PLIN2 and PLIN3 could increase the access of PNPLA2 to LD for lipolysis [34]. The downregulation of *Pnpla3* during fasting [35] may increase the availability of ABHD5 to activate PNPLA2 independently from PLIN5. 17beta HSD13 regulates PNPLA2-ABHD5 interaction as well in a PKA-dependent manner [36]. Second, PGC1a/PPARa can be activated by mechanisms independent from FA availability and hepatic lipolysis. NAD^+^ and AMPK can activate PGC1a/PPARa and support the expression of target genes during fasting in the liver [37]. While PNPLA2 dependent lipolysis is known to increase deacetylase activity of SIRT1 and activate PGC1a [16], SIRT1 can be directly phosphorylated by PKA as well [38]. PPARa expression during fasting is upregulated by glucocorticoid that may play a dominant role in increasing transcription activity of PPARa [18]. It also is plausible that other phosphorylation sites of PLIN5 compensate for the loss of S155 phosphorylation and maintain lipolysis and nuclear translocation. In the study of recombinant PLIN5 proteins, S161 and S163 were shown to be phosphorylated in addition to S155 [11]. When S155A PLIN5 is overexpressed in the heart, there was an increase in S17 and S292 phosphorylation that may compensate for the loss of S155 phosphorylation [8]. However, significant reduction of phosphorylated PLIN5 in the liver of SA/SA mice in Phos-Tag gel supports that S155 is the predominant site being phosphorylated in PLIN5 in vivo. In agreement, [^32^P] incorporation into recombinant PLIN5 was reduced in S155A mutant but not in S161A or S163A mutants incubated with catalytic subunit of PKA [11].

While PKA dependent phosphorylation of PLIN5 is well established in cultured cell models, little is known for the extent by which PLIN5 phosphorylation changes in vivo when PKA is activated or PLIN5 expression level is increased physiologically. We utilized Phos-tag gel as a simple method to determine the proportion of phosphorylated PLIN5 in tissue protein lysates. Phos-tag gel of total liver lysate showed that the proportion of p-PLIN5/non p-PLIN5 did not change when PLIN5 increases in the liver upon fasting or leptin deficiency indicating that there is an increase in both phosphorylated and non-phosphorylated forms of PLIN5. In the heart, exercise significantly increased the proportion of p-PLIN5 indicating that the extent of PLIN5 phosphorylation differs dependent on tissues and context. As one proposed function of PLIN5 phosphorylation is to increase lipolysis to provide FA for FAO, the precise regulation of FA substrate is likely more critical in the heart that primarily depends on FAO for energy production leading to more dynamic regulation of PLIN5 phosphorylation. The difference in the proportion of phosphorylated PLIN5 may account for the extent and the direction of correlation between PLIN5 levels and PPARa target genes/mitochondrial function/FAO in different tissues. PLIN5 levels in the heart negatively correlates with mitochondria function, FAO, PPATA/PGC1a target genes [39; 40], while the positive correlation is reported for the skeletal muscle [41].

IRS2 was reduced both at mRNA and protein levels in the liver of fasted SA/SA male mice. IRS2 is considered to mediate insulin signaling primarily during fasting while IRS1 plays a predominant role during refeeding [42]. Transcriptional regulation plays an important role in increasing hepatic IRS2 expression during fasting. FOXO regulates IRS2 expression to create a feedback loop to support insulin signaling during fasting [43]. Glucagon increases CREB/CRTC2 complex and increases IRS2 expression for which PGC1a appears to be required [43; 44]. It requires further studies to determine whether p-PLIN5 interacts with known regulators of hepatic IRS2 expression or acts by a new mechanism during fasting in male mice.

Lipid and glucose homeostasis of SA/SA knock-in mice in the current study showed a good agreement with mice in which SA PLIN5 was overexpressed in the liver using adenovirus associated virus (AAV) in PLIN5 knockout mice [11]. Similar to SA/SA mice, liver TG contents, TG secretion, plasma TG, plasma FFA, and plasma beta hydroxybutyrate were not altered in AAV-SA PLIN5 expressing PLIN5 knockout mice compared with AAV-WT PLIN5 expressing mice [11]. Although lipolysis measured in liver slice ex vivo was lower in AAV-SA liver compared with AAV-WT liver, authors concluded that overall impact of phosphorylation resistant PLIN5 on hepatic lipid homeostasis is limited [11]. Interestingly, AAV-SA PLIN5 expressing PLIN5 knockout mice showed elevated blood glucose at oral GTT compared with AAV-WT PLIN5 expressing male mice, which was associated with lower glucose-stimulated insulin secretion [11]. 1-year-old SA/SA male mice in the current study showed mild elevation in fasting glucose and elevated glucose at ip GTT performed after overnight fasting. As ITT did not differ between SA/SA and WT mice, it is plausible that glucose intolerance of SA/SA male mice is due to impaired insulin secretion. However, serum insulin level was not reduced when fasting glucose was elevated in SA/SA mice in our study. Also, ITT was performed after short fasting to keep blood glucose levels at time 0, which might limit the effect of SA PLIN5. Further studies are required to determine whether reduced hepatic IRS2 level is sufficient to explain glucose intolerance or other mechanisms exist for the dysregulation of glucose homeostasis in SA/SA male mice.

The current study has several limitations. In LC-MS/MS analysis of semi-purified LD fraction of the liver, overnight fasting increased phosphorylation of PLIN5 peptide containing S155 but not that of non-phosphorylated peptide. In comparison, total liver lysate analyzed by Phos-tag gel increased both phosphorylated and non-phosphorylated PLIN5. Fasting increases PLIN5 in non-lipid droplet fraction and non p-PLIN5 may be preferentially distributed to non-LD fractions. Thus, further studies could address p-PLIN5/non p-PLIN5 in different fractions including the nuclear fraction. It also needs to be noted that RNA sequencing was performed using female mice since female mice are reported to show more changes in hepatic gene expression upon fasting [29]. It is possible that RNA sequencing of SA/SA male liver may yield additional genes that are differentially regulated. Nevertheless, qPCR and Western blot did not reveal the major impact of SA/SA PLIN5 mutation on previously characterized PGC1a/PPARa targets important for FAO and mitochondrial mass in the liver. Although the impact of SA/SA knock-in is limited in fasted liver, it remains to be determined whether p-PLIN5 is required in other tissues and in a different context, especially in a tissue like the heart where the proportion of p-PLIN5 can be dynamically regulated. While sexual dimorphism exists in many genes, it requires further studies to address why hepatic *Irs1* and *Irs2* expression was differentially regulated between male and female SA/SA mice.

## 5. Conclusion

Despite p-PLIN5’s activity as a nuclear transcriptional regulator in cultured cells, PLIN5 SA/SA knock-in mice indicated that S155 phosphorylation is dispensable for the upregulation of lipid metabolism genes during the fasted state in vivo. Impairing phosphorylation also had little effect on serum lipids or liver TG. However, SA/SA male mice showed decreased IRS2 expression in the liver, which may explain glucose intolerance in SA/SA male mice.

## CRedit authorship contribution statement

**Corinne E Bovee:** Writing – original draft, data curation, formal analysis, validation, visualization, writing – review and editing. **Ryan P Grandgenett:** Data curation, formal analysis, validation, visualization, writing – review and editing. **Michelle Trevino:** Conceptualization, data curation, formal analysis, methodology, visualization, writing – original draft, writing – review and editing. **Sucharita Dutta:** Data curation, formal analysis, methodology, writing – original draft, writing – review and editing. **Spencer J Peachee:** Data curation, writing – review and editing. **Shayla Kopriva:** Data curation, writing – review and editing. **Farakh Haider:** Data curation, writing – review and editing. **Siming Liu:** Methodology, Supervision, writing – review and editing. **Gourav Bhardwaj:** Data curation, writing – review and editing. **Christie Penniman:** Data curation, writing – review and editing **Brian T, O’Neill:** Formal analysis, resources, methodology, supervision. **Yumi Imai:** Writing – original draft, conceptualization, formal analysis, funding acquisition, project administration, resources, supervision, validation, writing – review and editing.

## Acknowledgement

Data presented herein were obtained at the Genomics Division of the Iowa Institute of Human Genetics which is supported, in part, by the University of Iowa Carver College of Medicine. YI is supported by the National Institutes of Health (R01-DK090490), US Department of Veteran Affairs Biomedical Laboratory R&D (BLRD) (I01 BX005107), and Fraternal Orders of Eagles Diabetes Research Center at the University of Iowa. BTO is supported by VA Merit Review Award Number lO1 BXOO4468 from the US Department of Veterans Affairs R&D (BLRD) Service. Mass spectrometric analysis was performed at Leroy T. Canoles Cancer Research Center, Eastern Virginia Medical School. We thank Dr. Tina Tootle, Dr. Michelle Giedt, and Mr. Israel Wipf of University of Iowa for helpful discussion.

## Data availability

Data will be made available on request to the corresponding author.

## Abbreviation

LD: Lipid droplet
PLIN1-5: Perilipin 1-5
TG: Triglycerides
PKA: Protein kinase A
FA: Fatty acids
FAO: Fatty acid oxidation
PNPLA2: Adipose triglyceride lipase
ABHD5: abhydrolase domain containing 5
PGC1a: peroxisome proliferator-activated receptor-gamma coactivator
SIRT1: Sirtuin 1
PPARa: peroxisome proliferator-activated receptor alpha
IRS2: insulin receptor substrate 2
WT: Wild type
OA: Oleic acid
FBS: fetal bovine serum
RIN: RNA integrity number
GTT: glucose tolerance test
ITT: insulin tolerance test
SEM: standard error of mean
IRS1: Insulin receptor substrate 1
ASO: Antisense oligonucleotide
NEFA: Non-esterified fatty acids

