## Supplementary methods for "Perilipin 5 Phosphorylation is Dispensable for Upregulation of Hepatic Lipid Metabolism Genes upon Fasting but Required for Insulin Receptor Substrate 2 Expression in Male Mice"

### *1. Genotyping of PLIN5 S155A nice:*

Restriction enzyme digest of PCR products encompassing PLIN5 S155A was used to distinguish between WT and mutant S155A mice. DNA was extracted from mouse ear tissue using Qiagen DNeasy kit (Qiagen, Valencia, CA) according to manufacturer instructions. PCR was used to amplify PLIN5 using primers with forward sequence (CTT GGA GGG AAT CAG GCA TGT) and reverse sequence (ACT TTC TGG GGT GCA TAG TGG). Invitrogen Platinum Hot Start PCR 2X master mix and GC Enhancer were added (Invitrogen). The manufacturer's protocol and recommended PCR conditions were followed. PCR product was run on 2% agarose gel to identify presence of single 564 bp band expected for PLIN5 amplifcation. The PCR product was then digested using ThermoFisher FastDigest Sacl in FastDigest buffer and water (ThermoFisher). The samples were incubated in 37 °C for 1 h and then ran on a 2% agarose gel. Following Sac1 digest, the S155A mutant allele appears as two bands at 240 and 124 bp, but the WT allele is not undigested and appears at 564 bp band only. The difference in band appearance was used to distinguish between WT, heterozygous (WT/SA) , and homozygous (SA/SA) mice.

### *2. Peptide/protein identifications by Liquid Chromatography Electrospray Ionization Tandem Mass Spectrometric (LC/ESI-MS/MS) analysis*

Mass spectrometric analysis was performed at Leroy T. Canoles Cancer Research Center, Eastern Virginia Medical School. The dried samples were dissolved with 20 µl of 0.1% formic acid in water. 2 µl each of sample was analyzed by LC/ESI-MS/MS using a Q-Exactive mass spectrometer (Thermo Fisher) with an Easy NanoLC-1000 system using data dependent acquisition with dynamic exclusion (DE=1) settings. The data dependent acquisition settings used were a top 12 higher energy collision induced dissociation (HCD) for the Q-Exactive MS. Resolving power for Q-Exactive was set at 70,000 for the full MS scan, and 17,500 for the MS/MS scan at m/z 200. LC/ESI-MS/MS analysis was conducted using a C18 column (75 µm x 150 mm). The mobile phases for the reverse phase chromatography were (A) 0.1% formic acid in water and (B) 0.1% formic acid in acetonitrile. A four-step, linear gradient was used for the LC separation (2% to 30% B in the first 47 min, followed by 80% B in the next 1 min and holding at 80% B for 12 min).

The Sequest algorithm was used to identify peptides from the resulting MS/MS spectra by searching against the combined Mouse protein database (a total of 69,036 sequences) extracted from Swissprot (version 57) with taxonomy "Mus Musculus" using Proteome Discoverer (version 1.3, Thermo Scientific). Searching parameters for parent and fragment ion tolerances were set at 15 ppm and 60 milli mass unit (mmu) for the Q Exactive MS. Other parameters used were a fixed modification of carbamidomethylation –Cys, variable modifications of phosphorylation (Ser, Thr, Tyr), and oxidation (Met). Trypsin was set as the protease with a maximum of 2 missed cleavages. Raw files were searched against the five isoforms of Perilipin protein sequence (along with 500 other random proteins and reversed proteins as decoys) using Byonic [1] with a peptide tolerance of 15 ppm; an MS/MS tolerance of 20 ppm for HCD data; the carbamidomethylated cysteine as fixed modification and oxidation of Met and phosphorylation of Ser, Thr, and Tyr as variable modifications. Byonic scoring gives an indication of whether modifications are confidently localized.

#### *3. Bulk RNA-sequence of the liver*

Sequencing libraries was prepared from 500 ng of DNase I-treated total RNA using the Illumina TruSeq stranded mRNA library preparation kit (Cat. #RS-122-2101, Illumina, Inc., San Diego,

CA). The molar concentrations of the resulting indexed libraries were measured using the 2100 Agilent Bioanalyzer (Agilent Technologies) and combined equally into a pool for sequencing. The concentrations of the library pools were measured using the Illumina Library Quantification Kit (KAPA Biosystems, Wilmington, MA) and sequenced on the Illumina NovaSeq 6000 genome sequencer using 100 bp paired-end SBS chemistry.

Barcoded samples were pooled and sequenced using an Illumina NovaSeq 6000 in the Iowa Institute of Human Genetics Genomics Core Facility. Paired-end reads were demultiplexed and converted from the native Illumina BCL format to FASTQ format using a custom python workflow wrapper to Illumina's 'bcl2fastq' conversion utility. FASTQ data were processed with nf-core/rnaseq (v3.12), a best-practices pipeline available at the open-source 'nf-core' project (https://nf-co.re, Nextflow version 22.10). Reads from the samples were aligned against the mouse reference genome 'GRCm38' using the STAR aligner [2] and quantified with 'salmon' [3]. Samtools was used in conjunction with Qualimap and MultiQC to inspect alignment results [4-6]. Length-normalized gene-level counts from the STAR/salmon pipeline were used for differential gene expression analysis with DESeq2 [7]. Bioconductor package 'PCAExplorer' was used for exploratory analysis [8]. The DE gene lists were analyzed using AdvaitaBio's iPathwayGuide (https://www.advaitabio.com/ipathwayguide/). This software analysis tool implements the 'Impact Analysis' approach that takes into consideration the direction and type of all signals on a pathway [9]. The raw FASTQ files and associated metadata have been made available for download at GEO accession GSE275608.

**Supplementary Table 1. Antibodies used for the study**

| | Dilution | Source |
|---|---|---|
| guinea pig anti-PLIN5 for Fig. 1 | 1:1,000 | GP31 Progen Biotechnik |
| guinea pig anti-PLIN5 for all others | 1:5,000 | ProSci (custom made#) |
| rabbit anti-phospho-PKA substrate (RRXS*/T*) | 1:1,000 | 9624 Cell Signaling Technology |
| rabbit anti-GAPDH | 1:1,000 | 2118, Cell Signaling Technology |
| rabbit anti-IRS2 | 1:1000 | 4502, Cell Signaling Technology |
| rabbit anti-IRS1 | 1:1000 | 2382, Cell Signaling Technology |
| Total OXPHOS Rodent WB Antibody Cocktail | 1:1000 | ab110413, Abcam |
| 800 CW donkey anti-Guinea pig IgG | 1:10,000 | 925-32411 LI-COR Biosciences |
| HRP conjugated goat anti guineapig IgG | 1:10,000 | A7298 Millipore Sigma |
| HRP conjugated goat anti rabbit IgG antibody | 1:2000 | 12-348 Sigma Aldrich |

#: C terminal aa 451-463 of human PLIN5 peptide was used as immunogen.

**Supplementary Figure. 1**

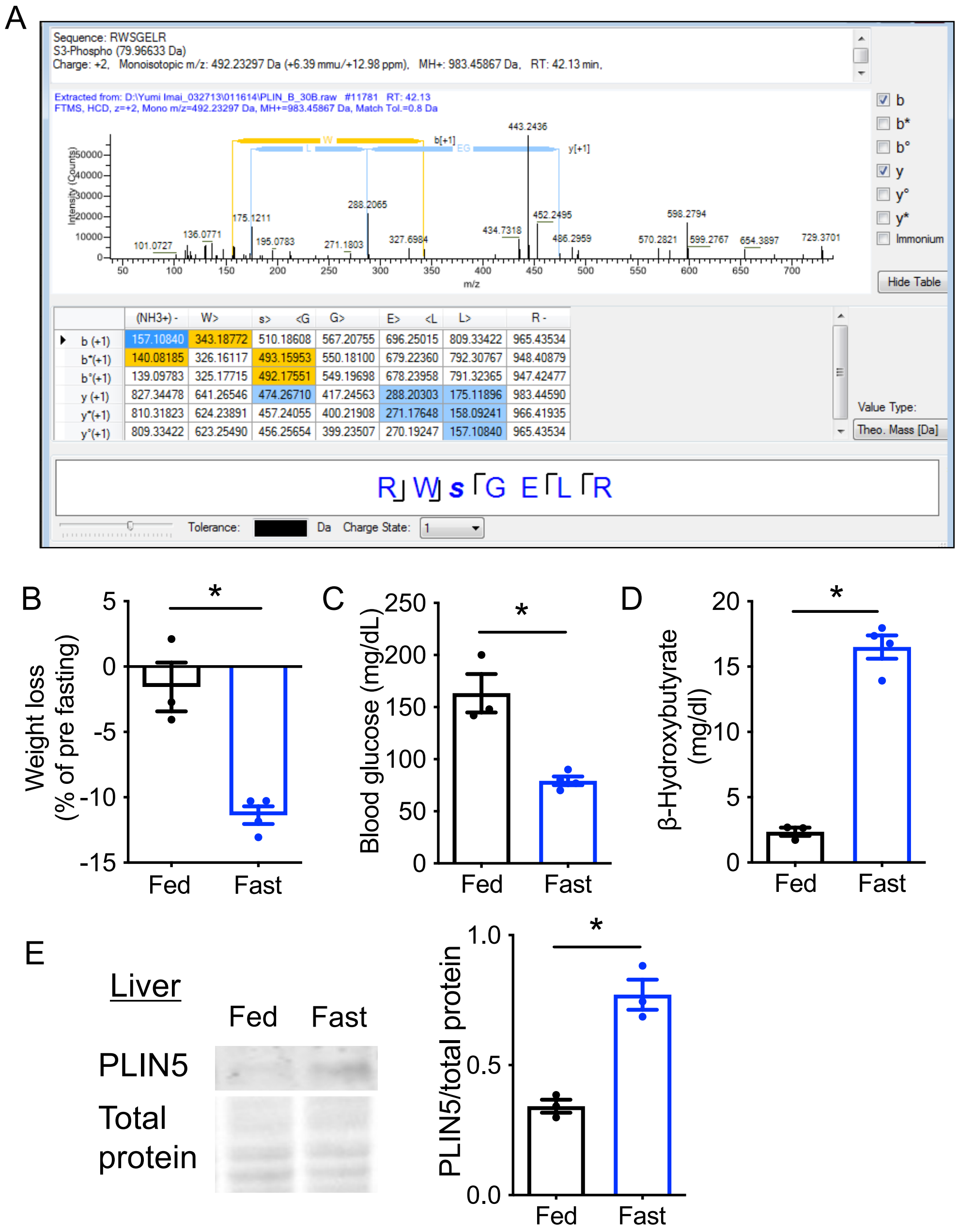


(A) Peptide consensus view denoting b and y ion masses corresponding to PLIN5 peptide RWSGELR containing S155 obtained from immunoprecipitation of PLIN5 from primary hepatocytes overexpressing WT PLIN5 (Fig. 1A). (B-D) Metabolic parameters including (B) body weight loss, (C) blood glucose levels, and (D) serum β-hydroxybutyrate measured in male mice fed *ad libitum* or fasted overnight. n = 3-4. (E) Western blot of PLIN5 in liver from fed and fasted mice using total protein staining as an internal loading control. n = 3. Means ± SEM; *$p < 0.05$ by Student’s t test.

**Supplementary Figure. 2**

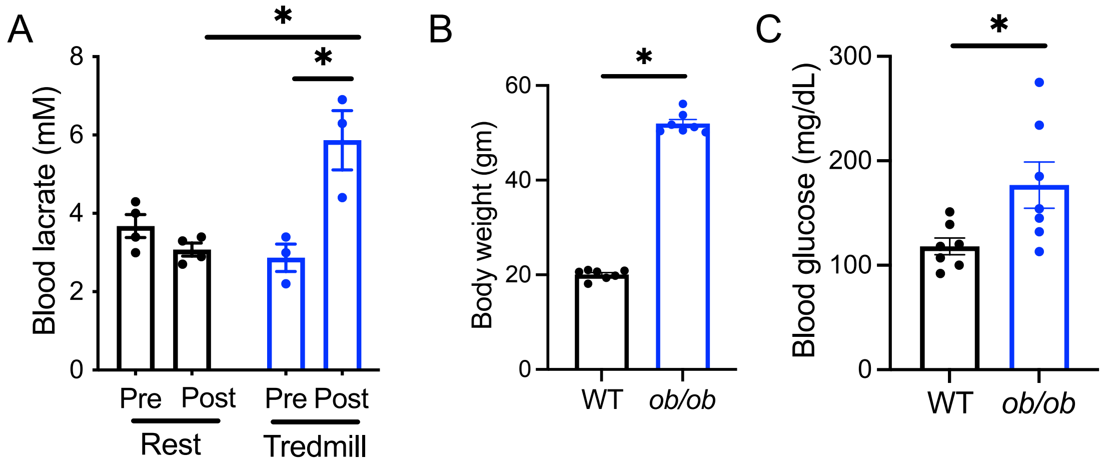


(A) Blood lactate levels pre and post treadmill in 3-month-old male WT mice. Control (rest) was not run on treadmill. n=3-4. 2-way ANOVA; p<0.05. *p<0.05 by Tukey's multiple comparison test. (B) Body weight and (C) blood glucose of WT and *ob/ob* female mice. n=7. Means ± SEM. *$p < 0.05$ by Student's t test.

**Supplementary Figure. 3**

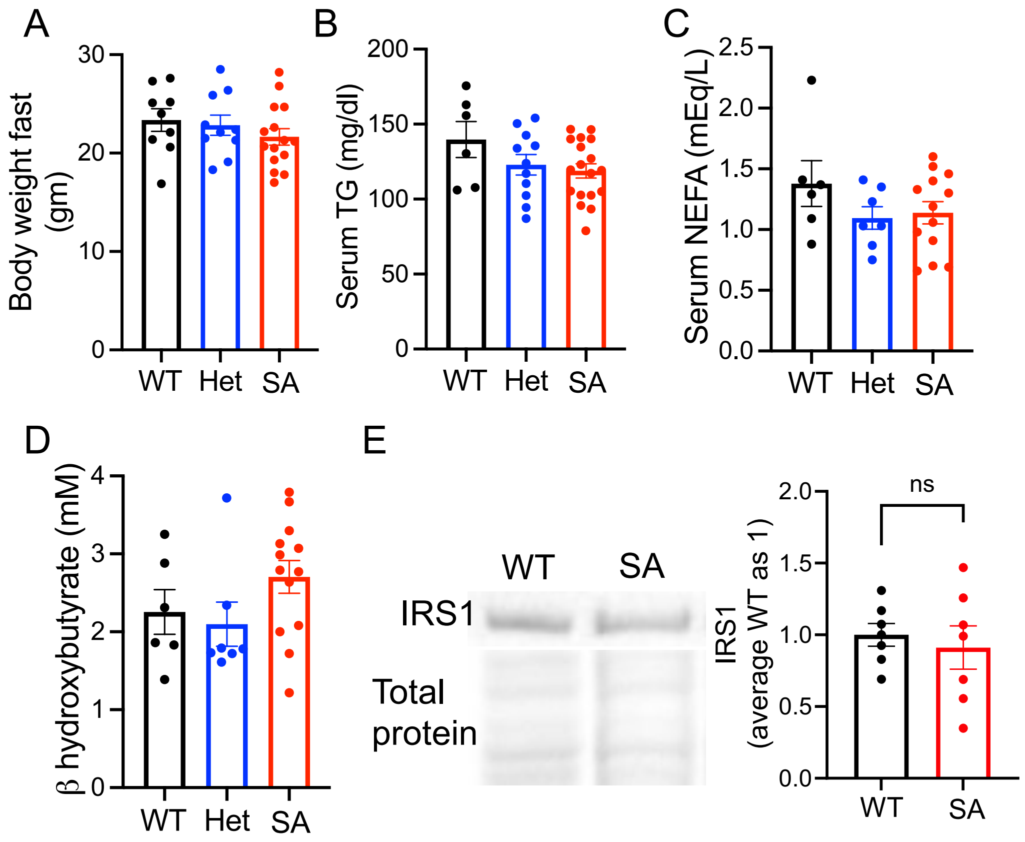


(A) Fasting body weight, (B) serum TG, (C) serum non-esterified fatty acids (NEFA), and (D) serum β-hydroxybutyrate of 4-month-old female mice. Each dot represents one mouse. n= 6-9 (WT), 7-11 (het), and 13-18 (SA). (E) Representative blot and densitometry of IRS1 normalized for total protein stain in the liver of overnight fasted female mice. n=7. Means ± SEM.

**Supplementary Figure. 4**

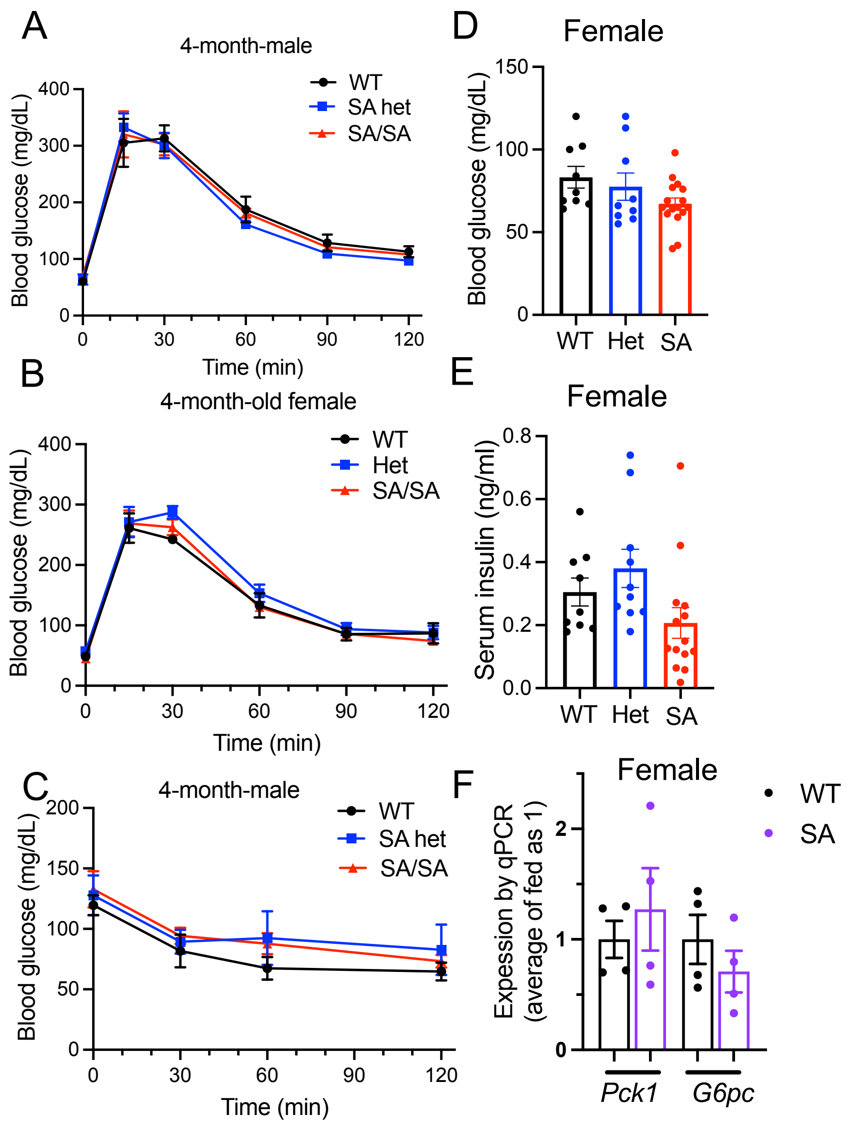


(A-B) i.p glucose tolerance test compared WT, SA het (het), and SA/SA (SA) male (A) and female (B) mice at 4-month-old male. Males, n= 5 (WT), 3 (het), and 4 (SA). Females, n= 3 (WT), 4 (het), and 7 (SA). (C) i.p insulin tolerance test compared male WT, het, and SA mice at 4-month-old. n= 5 (WT), 3 (het), and 4 (SA). (D) Blood glucose and (E) serum insulin of 4-month-old female mice fasting overnight. Each dot represents one mouse. n= 9 (WT), 9-10 (het), and 14-16 (SA). (F) qPCR compared the expression of genes for gluconeogenesis in the liver of 6-month-old WT vs SA female mice fasted overnight. n=4. Mean ± SEM. *$p<0.05$ by Student's t test.

**Supplementary Figure. 5**

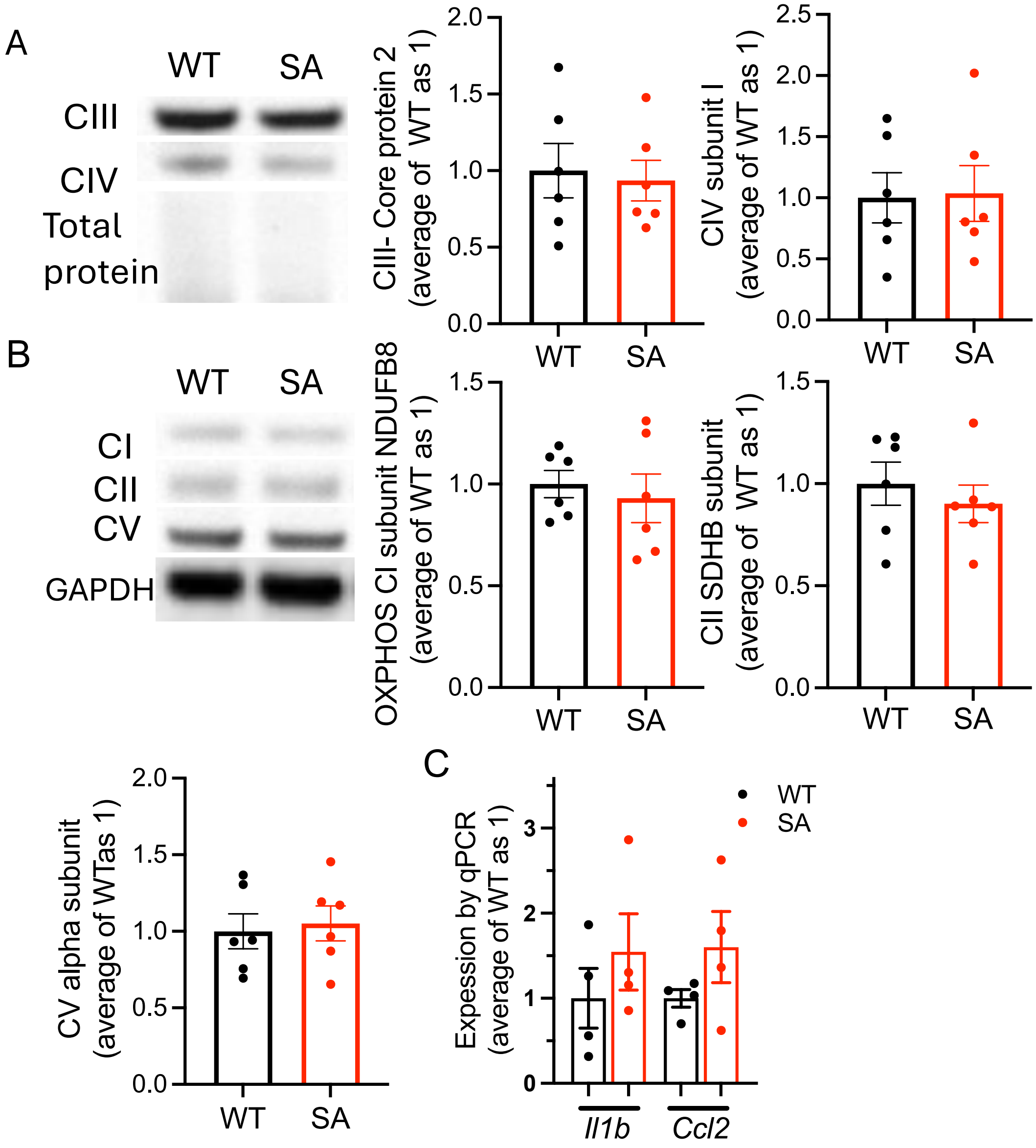


(A, B) Representative Western blot and densitometry of (A) OXPHOS complex III and V and (B) I, II, and IV in the liver of fasted male WT and SA mice. n=6. (C) qPCR comparing the expression of inflammatory genes in the liver of fasted male WT and SA mice. n=4. Mean ± SEM.
